# Atypical RanGAP drives nucleocytoplasmic transport in a parasitic Alveolate

**DOI:** 10.64898/2026.08.05.743068

**Authors:** Pravin S. Dewangan, Michael L. Reese

**Affiliations:** Department of Pharmacology, University of Texas, Southwestern Medical Center, Dallas, TX USA; Department of Biochemistry, University of Texas, Southwestern Medical Center, Dallas, TX USA

## Abstract

Transport of macromolecules between the nucleus and cytoplasm requires a gradient of the small GTPase, Ran. Ran:GTP marks the nucleus because Ran activity requires a cytoplasmic GTPase activating protein (RanGAP) for GTP hydrolysis. As expected for such central and essential cellular machinery, both Ran and RanGAP are conserved across the vast majority of eukaryotes. Many alveolates, including apicomplexan parasites, however, lack a canonical RanGAP. Here, we biochemically purify RanGAP activity from the model alveolate *Toxoplasma gondii*. We demonstrate that this activity is provided by a neofunctionalized RabGAP-fold protein, called TBC9. We find that TBC9 is sufficient to provide RanGAP activity in yeast, and is absolutely required to maintain active nucleocytoplasmic transport in *Toxoplasma*. By purifying *Toxoplasma* Ran and TBC9, we demonstrate that TBC9 is a robust and specific GAP for Ran. Finally, we delineate an essential, conserved low complexity motif in the C-terminus of TBC9 that drives interaction with Ran and is required for its full RanGAP activity. We use this C-terminal motif to identify TBC9 orthologs in all non-ciliate clades of Alveolata, which suggests that the canonical RanGAP has been replaced by the neofunctionalized TBC9 RanGAP in these lineages.

## Introduction

One of the hallmarks of a eukaryotic cell is the regulated transport of macromolecules between the nucleus and cytoplasm (Figure 1A). Because this process is thought to have arisen early in eukaryote evolution, the proteins that drive it are generally well conserved and easily identifiable across eukaryote diversity (1–5). The nuclear pore complex acts as a gateway through which nucleocytoplasmic transport (NCT) is regulated (4, 6–12). Cargo is carried through the pore by interacting with soluble transport proteins called importins and exportins (4, 6, 7). Fundamental to this process is the maintenance of the gradient of a small GTPase called Ran, which is exclusively GDP-bound in the cytoplasm and GTP-bound in the nucleus (13). Importins and exportins have high affinity for Ran:GTP, but low affinity for Ran:GDP, and binding and release of Ran regulates the binding and release of the trafficked cargo. In the nucleus, Ran:GDP is quickly converted to Ran:GTP by its guanine exchange factor (GEF), RCC1 (14–16). Ran:GTP binding to exportins in the nucleus triggers release of the trafficked cargo. Nuclear importins and exportins also must bind Ran:GTP to traffic to the cytosol (8, 17, 18). Once in the cytosol, the Ran GTPase activating protein (RanGAP) triggers GTP hydrolysis. This allows Ran (and export cargo) release by its export partners and Ran shuttle back to the nucleus (Figure 1A) (7, 8, 19–21).

**Figure 1:**
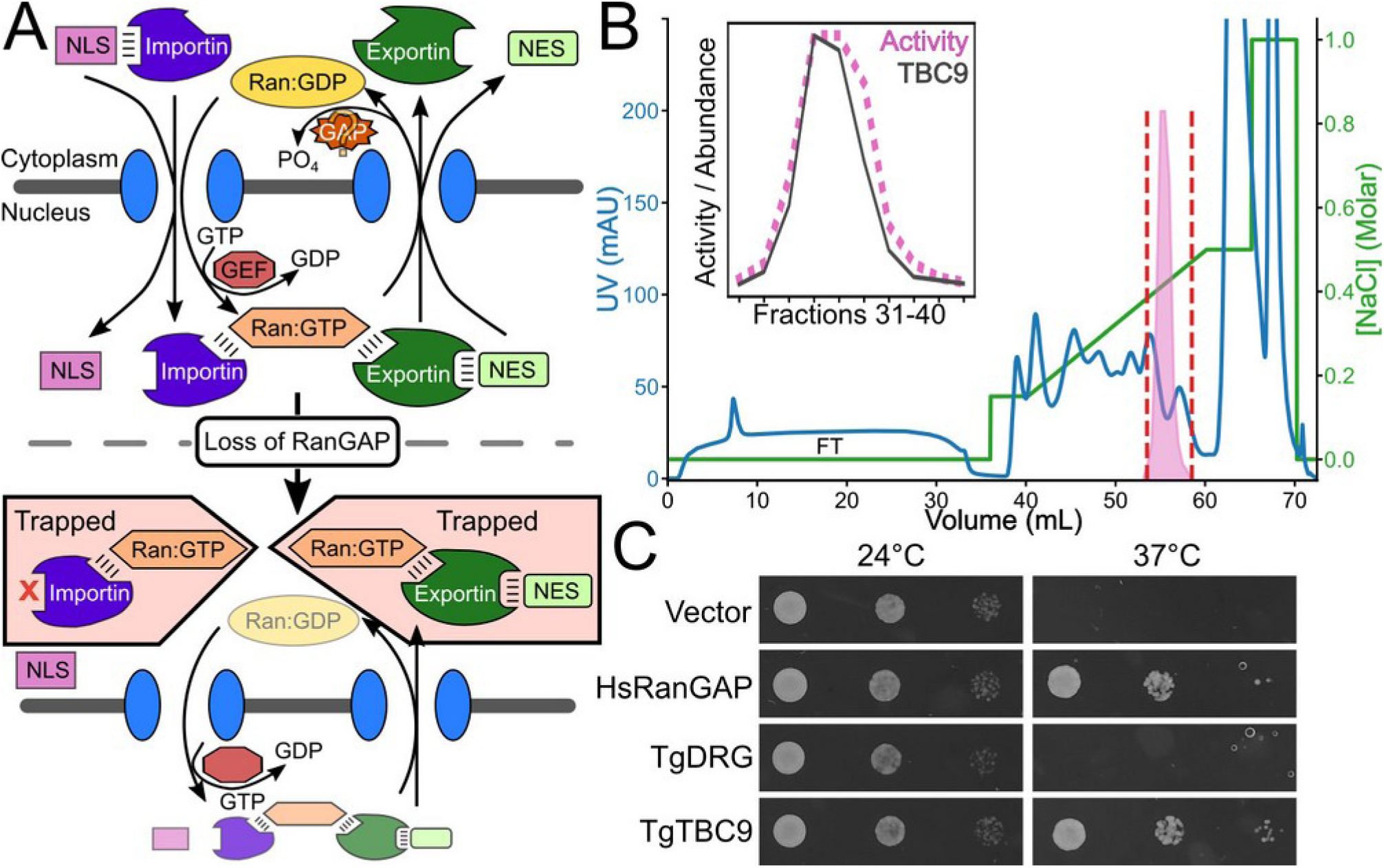
Purification of RanGAP activity from *Toxoplasma* lysate. (A) Cartoon overview of major proteins regulating nucleocytoplasmic transport. When RanGAP function is disrupted, Ran:GTP builds up in the cytosol, trapping exportins, importins, and their cargoes. (B) Anion exchange chromatogram showing fractions taken for down-stream MS overlayed with RanGAP activity (pink peak). Inset: correlation between RanGAP activity and TgTBC9 abundance in these fractions. (C) *rna1-1* yeast (RanGAP temperature-sensitive) expressing the indicated proteins grown at permissive (24°C) and restrictive (37°C) temperatures.

While the importins, exportins, Ran, and its GEF, RCC1, and the RanGAP are well conserved throughout most of eukaryotes, a subset of alveolates appear to be missing the canonical RanGAP protein (22, 23). There is a diversity of folds that can provide GAP activity, and all known GAP domains for small GTPases bind the GTPase active site. Notably, all RanGAPs identified to date (*e.g.* from opisthokonts, green plants, and Excavata) have a core leucine-rich-repeat (LRR) GAP domain (24–26). While ciliates like *Tetrahymena* encode a predicted LRR-family RanGAP, there is no identifiable RanGAP encoded in the genomes of sequenced Apicomplexa, Chromerids, Perkinsozoa, or dinoflagellates (which comprise the major alveolate phyla other than Ciliates; Figure S1, SI Data S1-4)(23).

We have purified RanGAP activity from the model Alveolate *Toxoplasma gondii*, which we identified as provided by the essential RabGAP-fold protein, TBC9. We demonstrated that TBC9 acts as a RanGAP in both heterologous and native cellular systems, and is sufficient to robustly activate Ran *in vitro*. Furthermore, we delineated a conserved low complexity C-terminal region of the alveolate TBC9 that differentiates it from other non-RanGAP encoding TBC domains. We show that this region assists in binding Ran, and highlights how a simple low complexity sequence may assist in the evolution of new protein activities.

## Results

### Toxoplasma Ran GTPase requires a non-canonical GAP for full activity

While a previous study suggested that apicomplexan parasites, including *Toxoplasma*, lack a canonical leucine-rich repeat (LRR)-RanGAP (22, 23), it remained possible that the *Toxoplasma* RanGAP had merely diverged sufficiently that it was difficult to identify with tools such as BLAST. To address this, we created a master alignment of all LRR proteins in *Toxoplasma*, baker’s yeast, the red algae *Chondrus crispus*, and RanGAPs from diverse organisms, and used IQTREE (27) to estimate a phylogeny (Figure S1). Consistent with the previous study and databases such as OrthoMCL (28), we found no *Toxoplasma* LRR proteins that clustered with known RanGAPs.

It remained possible that the *Toxoplasma* Ran (TgRan; TG*_248340) had higher intrinsic activity than the protein from other organisms and therefore did not require a GAP to turn over GTP. To test this, we recombinantly expressed and purified TgRan and compared its ability to hydrolyze GTP in the presence or absence of purified human RanGAP protein. We found that, like most other small GTPases, TgRan has very low intrinsic activity (Figure S2A). In addition, we found that TgRan was modestly activated by the human RanGAP (Figure S2A). Taken together, these data demonstrate that *Toxoplasma* must encode a protein that is evolutionary distinct from LRR-RanGAPs to maintain the Ran gradient in cells.

### Purification of RanGAP activity from Toxoplasma lysate identifies candidate Ran activating proteins

RanGAP was originally identified through temperature sensitive screens in yeast (29, 30) and biochemical fractionation of the human protein by following RanGAP activity (31, 32). To identify the *Toxoplasma* RanGAP, we opted to adapt the protocol used to identify the human protein by enriching RanGAP activity from cell lysate. Using an available high-throughput GTPase assay coupled to luciferase activity (33), we first verified that we could measure RanGAP activity in cell lysate of parasites that had been filtered and washed to remove host cell contamination. We found that *Toxoplasma* lysate has >10-fold higher GAP activity towards TgRan per protein equivalent than human lysate (Figure S2B). In addition, we confirmed that there was minimal contamination of the human LRR-RanGAP in our preparation (Figure S2C). We identified RanGAP activity in both the soluble and membrane (extracted with Triton-X100) fractions of the parasite lysate. To reduce the sample complexity, we chose to follow only the soluble activity. To enrich RanGAP activity in the lysate, we took the soluble protein after a 40% ammonium sulfate cut, dialyzed it, and further separated it by anion exchange chromatography. Because mass spectrometry is able to identify proteins from much more complex mixtures than the N-terminal sequencing used historically, we reasoned we did not have to purify the *Toxoplasma* RanGAP activity to homogeneity. Instead, we used isobaric tagging to allow us to quantify the relative amounts of each protein in the 10 fractions of our final chromatography with RanGAP activity (Figure 1B). We then measured the covariance between biochemical activity and individual protein abundance in each sample using Pearson’s correlation (PCC).

While our enriched sample was still quite complex (>1000 proteins identified by mass spectrometry), only 14 proteins had abundance that covaried with RanGAP activity (PCC ≥ 0.95; SI Data S5). Many of these proteins, including our top hit (DRG GTPase; TG*_233260) appeared to be contaminants from a complex that regulates protein translation. Intriguingly, our second highest hit, TBC9 (PCC=0.98; TG*_226950), is a ∼36 kDa protein with a RabGAP TBC (Tre2-Bub2-Cdc16) fold. TBC9 was recently identified as the only essential TBC-domain containing protein in the parasite, and has been suggested to activate multiple *Toxoplasma* Rabs (34, 35). TBC domains, however, are not thought to interact with, or to regulate, Ran.

We next sought to quickly validate RanGAP function of our top candidates. To this end, we tested whether heterologous expression of either of our top two candidates could rescue the growth of the *rna1-1* (yeast RanGAP) temperature-sensitive mutant yeast (36) at restrictive temperature. As expected, we found that expression of the human LRR-RanGAP was able to rescue *rna1-1* growth, while a vector control was not (Figure 1C). Expression of DRG did not rescue *rna1-1*. We were excited to find, however, that expression of TBC9 was able to rescue *rna1-1* as robustly as the human LRR-RanGAP, indicating that TBC9 is sufficient to provide RanGAP activity in a cellular context (Figure 1C).

### Loss of Toxoplasma TBC9 quickly disrupts nucleocytoplasmic transport

We next asked whether TBC9 was required for maintenance of the parasite’s Ran gradient and therefore for nucleocytoplasmic transport. We created a strain in which TBC9 was C-terminally tagged with an AID degron (37, 38) and a 3xHA epitope to allow us to both track the protein and control its degradation with the addition of auxin (TBC9^AID^). Consistent with previous reports (34, 35), we found that TBC9^AID^ appears to localize to diverse membrane structures, including the parasite ER (Figure 2A). Encouragingly, TBC9^AID^ is completely nuclear excluded, which would be required of a RanGAP to maintain high Ran:GTP only in the nucleus. Addition of the auxin IAA resulted in loss of TBC9^AID^ protein within 2-3 h (Figure S3A), and parasite replication was blocked upon growth in auxin (Figure 2B). Previous studies demonstrated that induced degradation of TBC9^AID^ resulted in numerous morphological changes after >13-24 h (34, 35). We reasoned that if TBC9 is, indeed, the protein responsible for converting Ran:GTP to Ran:GDP in the cytosol, we should observe accumulation of cytoplasmic Ran within minutes after TBC9^AID^ has been completely degraded (Figure 1A).

**Figure 2:**
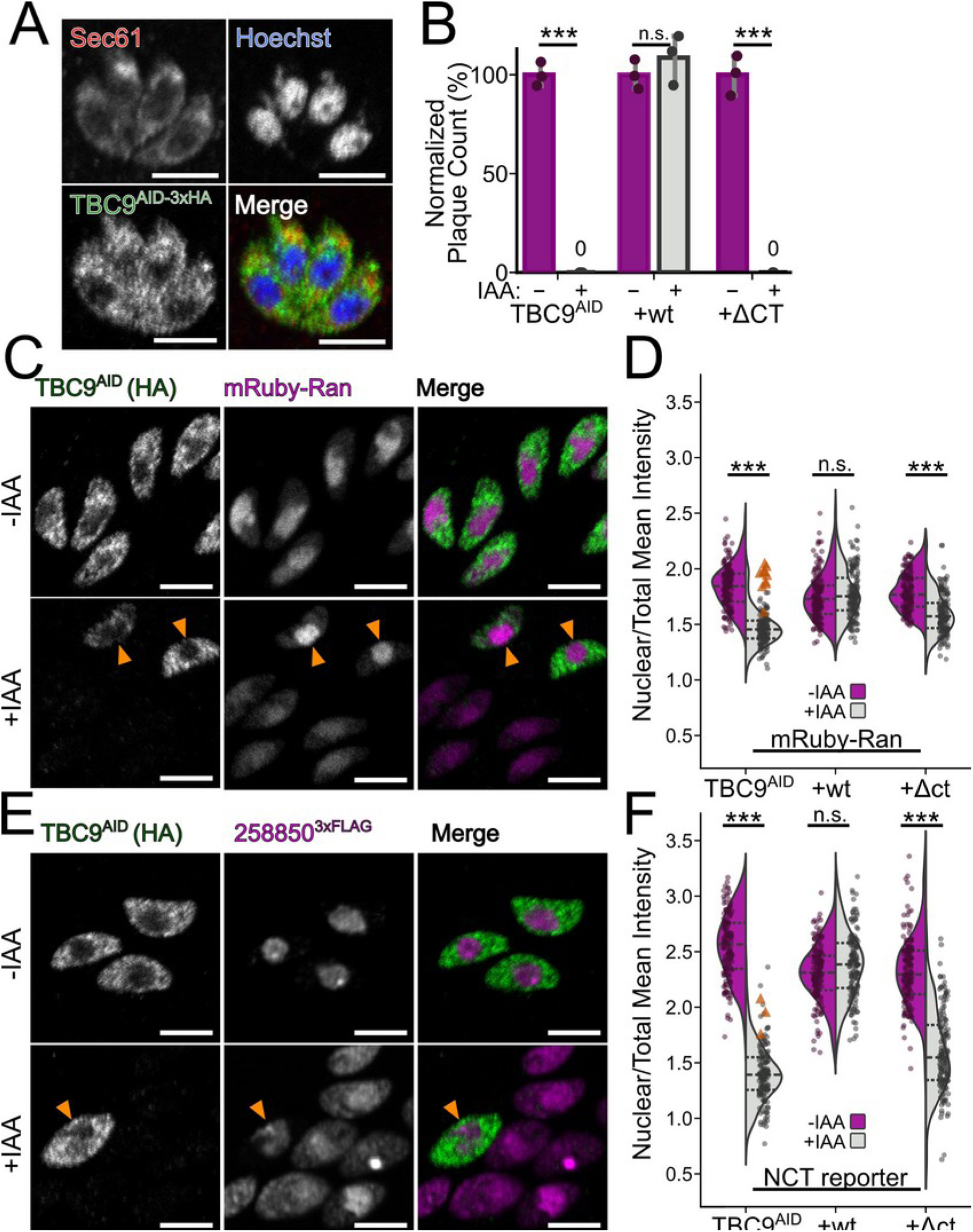
TgTBC9 is required for nucleocytoplasmic transport. (A) Confocal microscopy of TBC9^AID^ confirms punctate staining that is nuclear excluded and partially colocalizes with the ER marker Sec61. (B) TBC9^AID^ grown in +IAA are unable to form plaques. Growth is rescued by expression of a non-degradable wild-type copy but not a mutant lacking the conserved C-terminus. (C) Confocal micrograph showing 2h growth in +IAA media results in TBC9^AID^ parasites in which mRuby-Ran has begun to leak from the nucleus. Orange arrows indicate parasites in which TBC9^AID^ has not been degraded and maintain correct Ran localization. (D) Quantification of (C) n=100 parasites per n=3 biological replicates. ±25% confidence intervals are indicated. Orange arrows indicate parasites as in (C). (E) Confocal microscopy of the TGGT1_258850^3xFLAG^ NCT-reporter strain grown in ±IAA, annotated as in (C). (F) Data from (E) quantified as in (D). ***, p<0.001. p-values calculated by unpaired Student’s t-test with Benjamini-Hochberg correction. Scale bars: 5 μm.

To the observe how TgTBC9^AID^ degradation affects the Ran gradient, we created the TBC9^AID^ allele in parasites engineered to express Ran that had been endogenously tagged with an N-terminal mRuby3 (39) fusion. As expected, mRuby3-Ran is highly enriched in the nucleus in normal cells (Figure 2C). When parasites were incubated for 2 h in the presence of auxin, we found that mRuby3-Ran was significantly and visibly enriched in the cytosol (Figure 2C,D). Importantly, a minority of parasites grown in auxin still had some TBC9^AID^ protein, and in those cells, mRuby3-Ran was still concentrated in the nucleus (Figure 2C,D).

If the Ran gradient is disrupted, one would expect a simultaneous disruption of nucleocytoplasmic trafficking (Figure 1A) (40, 41). To test this, we created the TBC9^AID^ allele in parasites engineered to express the nuclear protein TGGT1_258850 (42) fused to 3xFLAG. As expected, we found that TGGT1_258850^3xFLAG^ was entirely nuclear-localized when parasites were grown in normal media (Figure 2E). Similar to our mRuby3-Ran reporter cells, we found that TGGT1_258850^3xFLAG^ began leaking from the nucleus into the cytosol quickly after degradation of TBC9^AID^ (Figure 2E,F). Notably, some cells had not completely degraded TgTBC9^AID^ after 2 h of incubation in auxin, and in those cells, TGGT1_258850^3xFLAG^ was still concentrated in the nucleus (Figure 2E,F). Together, these data lead us to argue that TBC9 is the *Toxoplasma* RanGAP and is absolutely required for maintenance of the Ran gradient and therefore nucleocytoplasmic transport.

### A conserved C-terminus in Toxoplasma TBC9 is essential for its function

TgTBC9 is a member of an ancient subfamily of TBC-domain proteins that are found in diverse protozoa, including trypanosomes (Figure 3A, SI Data S1,6-8; called “TBC-RootA” in a previous study (43)). Notably, while other apicomplexan TBC9 proteins can rescue TBC9 essentiality in *Toxoplasma,* the protein from *Trypanosoma brucei* cannot (34). This suggests that the alveolate TBC9 proteins may have a different function from other “TBC9-like” family members. We therefore tested whether trypanosome TBC9 could rescue RanGAP activity in yeast. Expression of trypanosome TBC9-like in *rna1-1* yeast failed to rescue growth at restrictive temperatures, consistent with a lack of RanGAP activity (Figure 3B). Notably, *T. brucei* encodes an essential canonical LRR-RanGAP (26), and therefore does not require an additional protein capable of RanGAP activity. We examined a sequence alignment of TBC domains from diverse organisms and identified a C-terminal helical extension and a motif in the last 15 residues of the protein that is highly conserved in apicomplexan and other myzozoan TBC9 proteins but missing in the majority of TBC9-like proteins outside of alveolates (Figure 3A, S4, SI Data S1,6-8). The only other TBC domains that contain this sequence are from the Haptophytes, distant relatives of Alveolates which likely share a plastid origin with Myzozoa (44–48). The C-terminal motif is highly acidic (9 of 15 residues Asp/Glu in TgTBC9), and is strikingly reminiscent of the conserved acidic C-terminus of Ran (Figure 3C,D). Intriguingly, the Ran C-terminus is thought to form regulatory interactions with a basic patch distant from its active site (49) and it is near this site that AlphaFold3 predicts the TBC9 C-terminus would sit (Figure 3E, Figure S4).

**Figure 3:**
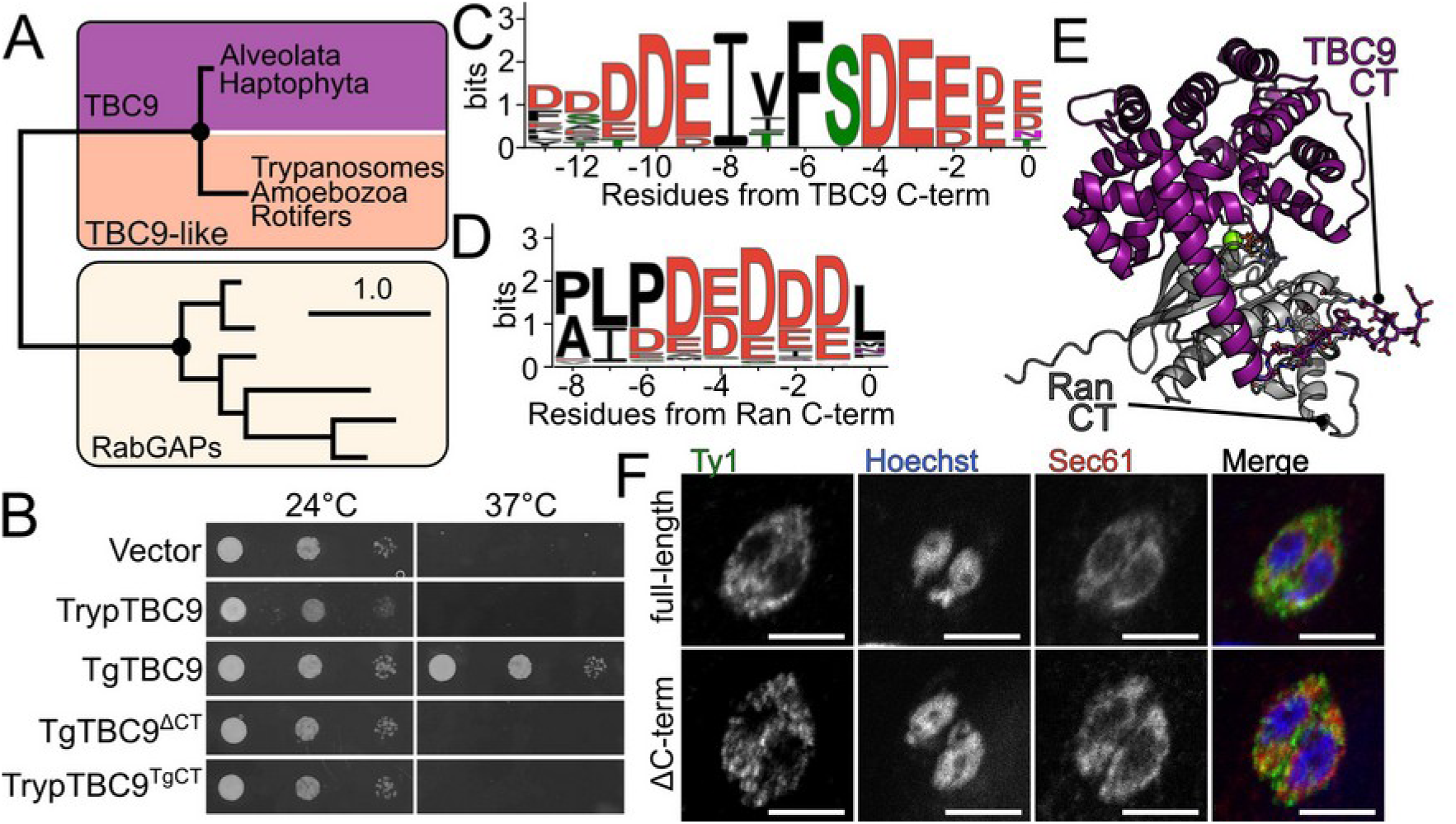
Alveolate TBC9 contains an essential conserved acidic C-terminal motif. (A) Cartoon phylogeny showing relationship of alveolate TBC9 with canonical TBC domains from yeast, human, and *Toxoplasma* versus TBC9-like domains from diverse protozoa. Black circles indicate >99% bootstrap support. Scale bar is 1 substitution per site. Complete phylogenetic tree and protein accessions are available in SI Data S1,S7, and guide with branch support in Figure S5. (B) *rna1-1* yeast expressing the indicated proteins grown at permissive (24°C) and restrictive (37°C) temperatures. Sequence logos of (C) the conserved alveolate TBC9 and (D) eukaryote Ran C-termini. (E) AlphaFold3 model of the Ran:TBC9 complex. Ran (gray), TBC9 (purple), GTP black, Mg ^2+^ green sphere. (F) Confocal microscopy comparing localization of TBC9^3xTy1^ and TBC9^ΔCT3xTy1^. Scale bars: 5 μm.

We therefore reasoned the TBC9 C-terminus may interact with the same region of Ran, thereby facilitating TBC9 function as a RanGAP. To test whether this region was required for TBC9 function in cells, we first tested whether TgTBC9^ΔCT^ (residues 1-300) could rescue the *rna1-1* yeast strain. We found that, unlike yeast expressing full-length TgTBC9 (residues 1-315), those expressing the mutant were unable to grow at a restrictive temperature (Figure 3B). We also found that a chimeric *T. brucei* TBC9 protein fused to the *Toxoplasma* C-terminus (residues 301-315) was still unable to rescue *rna1-1* growth (Figure 3B). Therefore, the TgTBC9 C-terminus is necessary, but not sufficient, for a TBC9- like protein to restore RanGAP function in these yeast. We next tested whether the C-terminus was required for TBC9 function in *Toxoplasma.* In the background of our TBC9^AID^ strains, we expressed an additional copy of either full-length wild-type TBC9^3xTy1^ or TBC9^ΔCT-3xTy1^ driven by the weak dhfr promoter. We found that both proteins showed similar punctate localization that, like TBC9^AID^, was nuclear excluded, suggesting that the TBC9 C-terminus is not required for nuclear exclusion (Figure 3F). Consistent with its lack of rescue in yeast, TBC9^ΔCT-3xTy1^ was unable to rescue plaque formation when parasites were grown in auxin, though the full-length protein fully restored plaque efficiency (Figure 2B). We next asked whether the TBC9 C-terminus was required for maintenance of the Ran gradient versus another, unknown function. We quantified the localization of mRuby-Ran and our nucleocytoplasmic reporter, TGGT1_258850^3xFLAG^, in the presence and absence of a brief 2 h auxin treatment. While the parasites expressing the full-length TBC9^3xTy1^ protein showed no disruption in either the Ran gradient or nucleocytoplasmic transport, those expressing TBC9^ΔCT-3xTy1^, exhibited phenotypes very similar to the parental TBC9^AID^ strains (Figure 2F). Notably, the TBC9^ΔCT-3xTy1^ phenotype was not as robust as that of the parental TBC9^AID^ strains, consistent with a partial, rather than complete, loss of RanGAP activity.

### Toxoplasma TBC9 is a robust and specific activator of Ran GTPase activity

We next sought to unambiguously determine TgTBC9 GAP activity against Ran versus Rabs. We first recombinantly expressed and purified *Toxoplasma* TBC9 and Ran. We purified Ran as quickly as possible, in a single day, to ensure we obtained a high percentage of GTP-bound protein. To quantify biochemical activity, we used a well-established continuous photometric assay (50, 51). By following single-turnover kinetics of TgRan:GTP, we would expect to fit TBC9 GAP activity to a pseudo-first order reaction (51). Comparing phosphate release from 20 μM TgRan ±TBC9 unequivocally demonstrated that TBC9 robustly activates TgRan (Figure 4A). However, we were unable to reliably fit the resulting data to an inverse exponential. Instead, the data fit well to a linear curve (pseudo-zeroth order), suggesting that the K_m_ of TBC9 for TgRan is much lower than the concentration at which we were conducting the assay (Figure 4A). Unfortunately, the assay sensitivity is such that we were unable to reduce the concentration of TgRan sufficiently to quantify kinetic parameters for the wild-type proteins.

**Figure 4:**
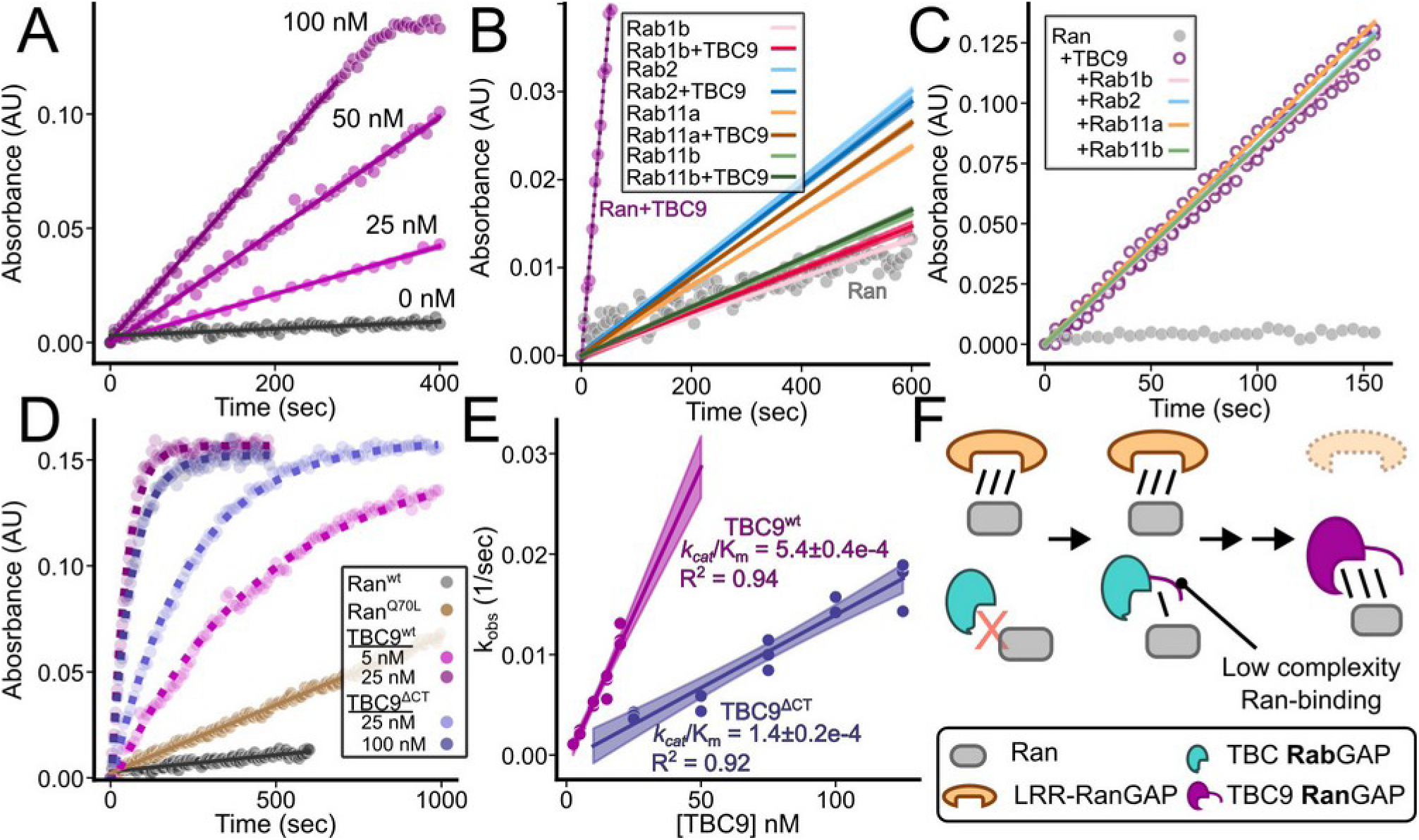
TBC9 is a strong GAP, specific for Ran. (A) Single turnover of 20 μM Ran:GTP (gray) with increasing amounts of TBC9 (purple) was observed by increase in free guanosine absorbance. (B) The indicated *Toxoplasma* GTPases (80 μM Rabs or 20 μM Ran) were incubated ±TBC9 (100 nM for Ran, 1 μM for Rabs). (C) Effect of the indicated Rabs on TBC9 RanGAP activity was measured by incubating 1 μM of the indicated Rabs (GTP-locked mutant) with 20 μM TgRan and 100 nM TBC9. (D) The indicated concentrations of TBC9^WT^ or TBC^ΔCT^ were incubated with 20 μM TgRan^Q70L^ and the resulting data fit to pseudo-first order kinetics (dotted lines). (E) kobs from fits from (D) graphed versus [TBC9] enables quantitative comparison of TBC9^wt^ versus mutant activities, with error expressed from confidence in fit. Shaded error for all fit curves is 95% ci. (F) Model by which low complexity sequence may have facilitated TBC9 neofunctionalization as a RanGAP.

Previous studies identified 4 *Toxoplasma* Rabs (Rab1b; TG*_214770, Rab2; TG*_312050, Rab11a; TG*_289680, and Rab11b; TG*_320480) that bound TBC9 in either a yeast-2-hybrid (34) or *in vitro* pull-down assay (35). While binding was observed, biochemical activity was never verified. We expressed and purified each of the four *Toxoplasma* Rabs that had been reported to interact with TBC9. We verified that the Rabs were GTP bound and had intrinsic activity (Figure S6A). We then measured the Rab GTPase activities in the presence and absence of TBC9 (Figure 4B). Surprisingly, incubation with TBC9 did not increase the activity of any of these Rabs, even at 10-fold higher concentration than used with TgRan (Figure 4B). Therefore, TBC9 is unable to function as a GAP for any of these Rabs. Both previous studies showed that TBC9:Rab interaction was increased in GTPase-dead mutants in the Rab switch loop II Gln (analogous to the TgRan^Q70L^ mutant used below). While such a mutant would not be expected to block activation by a TBC domain as it would by other GAPs (51), it remains possible that the Rabs bind TBC9 in a non-substrate orientation. We reasoned if that is the case, perhaps Rabs modulate TBC9 catalytic activity towards TgRan. We tested this by incubating 20 μM wild-type TgRan with 100 nM TBC9 and varying concentrations of Gln-mutant Rab. We were disappointed to find that none of the Rabs antagonized or augmented TBC9 activity against TgRan (Figure 4C). These data suggest that in evolving to act as a RanGAP, TBC9 has lost the ability to function as a RabGAP. In addition, any interaction TBC9 may have with Rabs appears to be independent of its biochemical activity as a RanGAP.

TBC domains are unusual in their mechanism of activation of GTPases in that they use a so-called “dual-finger” mechanism in which they provide both a catalytic Arg and Gln to complete the GTPase active site (51). Because of this, TBC domains often have the surprising ability to readily activate normally “GTP-locked” mutants in the GTPase switch-loop II Gln (Q70L in TgRan). Because the GTPase Gln interacts directly with the TBC domain to stabilize its binding (51), this mutation usually increases the K_m_ (reduced affinity). We therefore tested whether TBC9 could activate TgRan Q70L, reasoning that with the reduced affinity we may be able to obtain kinetic parameters. We recombinantly expressed and purified TgRan^Q70L^. Surprisingly, rather than being catalytically dead, TgRan^Q70L^ showed a ∼3-fold higher basal activity than wild-type, indicating that Gln70 is not required for hydrolysis in this protein (Figure 4D). This phenomenon has been previously observed in *Entamoeba* Rab21 (52), and suggests activation by a TBC domain may allow a relaxed evolutionary constraint at this residue in the GTPase.

We next tested TBC9 activity against TgRan^Q70L^. We found that, consistent with the known mechanism of TBC function, TBC9 was still able to robustly activate the mutant protein (Figure 4D). As expected, TBC9 has an apparently reduced affinity for TgRan^Q70L^, as we could now readily fit the data to a pseudo-first order reaction, allowing us to estimate kinetic parameters, albeit for the mutant protein (Figure 4D,E). We were also able to quantify TBC9^ΔCT^ activity, and found that the protein has ∼4-fold reduction in *k_cat_/*K_m_ for TgRan^Q70L^ as compared to full-length TBC9 (Figure 4D,E). Thus the TBC9 C-terminus is indeed required for full RanGAP activity. This is consistent with our cellular data that demonstrated TBC9^ΔCT^ is unable to rescue essentiality of TBC9 loss, though it exhibits an intermediate phenotype in nucleocytoplasmic transport (Figure 2). In the AlphaFold model of the Ran:TBC9 complex (Figure 3E), we found that the acidic residues in the TBC9 C-terminus are predicted to interact with a cluster of basic residues in Ran (Figure S6B). To further validate the AlphaFold model, we mutated the basic residues to Ala (R96A/K100A/R107A) in the background of TgRan^Q70L^. Consistent with the importance of this basic patch on Ran, full-length TBC9 showed a ∼2-fold reduction in activity towards the TgRan^R/K-Mut^ versus TgRan^Q70L^ (Figure S6C,D). If the TBC9 C-terminus does, indeed, interact with the basic patch, we would expect TBC9^ΔCT^ to have a *smaller* reduction in its activity towards TgRan^R/K-Mut^ than it does for the TgRan^Q70L^ protein. That is indeed, what we observed, as TgRan^R/K-Mut^ is a ∼2-fold poorer (rather than 4-fold) substrate for TBC9^ΔCT^ than for the full-length protein (Figure S6C,D). Therefore, TBC9 uses its C-terminus to recognize Ran through an interaction basic patch of residues distal from the Ran active site. However, this basic patch does not appear to provide 100% of the binding energy for the TBC9 C-terminus. There are an additional 6-7 Arg/Lys on Ran proximal to the TBC9 C-terminus in the model, and it is likely that the acidic motif binds this face in multiple conformations.

## Discussion

We have identified TBC9 as a neomorphic RanGAP that has replaced the canonical LRR protein in *Toxoplasma*. We demonstrated that TBC9 is sufficient to replace RanGAP function in a temperature-sensitive yeast strain, and necessary to maintain the Ran gradient in its native context. The effect on the Ran gradient, and on NCT, occurs immediately after TBC9 has been degraded. TBC9 loss-of-function has been previously demonstrated to cause major morphological changes in multiple organelles (34, 35), though these changes take >12 h to appear, and are likely secondary to disruption of NCT. We further demonstrated that TBC9 acts as a robust RanGAP *in vitro*, and has no apparent activity on other small GTPases with which it has been suggested to interact. Finally, we identified a conserved C-terminal motif in TBC9 that assists in binding Ran and is required for its full biochemical activity. While loss of the C-terminal motif shows only a modest ∼4-fold reduction in *in vitro* activity, expression of TBC9^ΔCT^ partially blunts the phenotype due to loss of the wild-type protein in cells, but is unable to complement its essentiality. This may be simply due to the requirements of tightly regulated nucleocytoplasmic transport. However, it is also possible that the TgTBC9 C-terminus has other moonlighting functions, as has been seen with the yeast RanGAP (53).

There are many folds that can serve as GAPs, and the only requirement seems to be recognition of a GTP-bound GTPase and stabilization of the transition state. Most GAP domains provide a so-called “Arg-finger” to complete the active site (54–56). The canonical LRR-RanGAP, however, makes no direct contacts with the Ran active site, so appears to act entirely through an allosteric mechanism (24). TBC domains function on the opposite side of the spectrum, providing not one, but two residues (Arg/Gln) to the substrate active site (51). This marked difference highlights potential variation in evolutionary pressure small GTPases activated by an LRR-RanGAP versus a TBC domain must be under. For example, both *Toxoplasma* Ran and *Entamoeba* Rab21 (52) do not require the switch-II Gln (Q70 in TgRan) for intrinsic activity, likely because TBC domains provide this residue. Intriguingly, the 4 TgRabs that have been demonstrated to bind TBC9 *in vitro* show improved binding when mutated at this site (34, 35). This is consistent with our data showing that these are not substrates. It is possible that TgTBC9 promiscuously binds Rabs outside of its active site, which would help explain its partitioning to membrane fraction (34) and its subcellular localization. Also, in spite of the additional binding surface provided by the TBC9 C-terminus, we found that Ran:TBC9, like the human Ran:LRR-RanGAP complex (24), is highly dynamic, even in the presence of a transition state analog (Figure S7). Understanding the structural and functional consequences of these interactions will require future studies.

Intriguingly, our bioinformatic analysis demonstrates that alveolates encode either an LRR-RanGAP (ciliates) or a TBC9 domain (Myzozoa; non-ciliate alveolates; Figure S8). This finding suggests that replacement of the LRR-RanGAP with TBC9 is widespread in Alveolates. Remarkably, we found that haptophytes, a clade of algae distantly related to alveolates, also appear to encode a TBC9 domain that contains the conserved alveolate-like C-terminus. Furthermore, as with Myzyzoa, we were unable to identify an LRR-RanGAP in available haptophyte genomes (SI Data S1-4). Intriguingly, Myzozoa and haptophytes are thought to share the evolutionary origin of their plastid (44, 45, 47). Nevertheless, while we identified TBC9-like domains in diverse protozoa, we were unable to identify these in any sequenced red algae. Notably, the *T. brucei* TBC9-like (or TBC-RootA), was found to co-immunoprecipitate with a large complex including Ran and the mRNA transporter Mex67 (57). Yet we found trypanosome TBC9 did not rescue the RanGAP-deficient *rna1-1* yeast, and *T. brucei* reportedly encodes an essential LRR-RanGAP (26). It remains an intriguing possibility, however, that TBC9-like domains represent an ancient clade (43) with varied functions in nuclear transport.

We were surprised to find that the low complexity region at the C-terminus of TBC9 is highly conserved in both length and content and essential for full TBC9 function both *in vitro* and in cells. That such a simple sequence can encode function suggests a potential mechanism by which TBC9 evolved RanGAP activity (Figure 4F). While low complexity regions are often overlooked, they may encode functional motifs sufficient to seed binding to a protein partner. We speculate that it was such binding that enabled TBC9 to replace the canonical RanGAP. This evolutionary trajectory was likely facilitated by polyploidy due to the proposed evolution of alveolates in which dueling red algal and protozoal genomes coexisted (47, 58). Regardless, our finding highlights the complexity of protein-protein interactions, and demonstrates how even simple sequences can encode important, yet hard to predict, functions.

## Materials and Methods

### PCR and Plasmid generation

The primers used in this study are listed in SI Data S9. All PCRs for construct generation used Phusion polymerase (New England Biolabs; NEB) and plasmids by Gibson assembly (NEB master mix). Deletion mutants were generated by enzyme inverse PCR. All plasmids were transformed into Mach1 (Invitrogen) competent *E. coli* for cloning and plasmid amplification.

### Cell culture and transfections

*Toxoplasma* tachyzoites were maintained on confluent monolayers of human foreskin fibroblasts (HFFs). HFFs were derived from deidentified tissue obtained from UTSW, and work was conducted under an institutional IRB waiver by the UTSW Human Research Protection Program. The HFFs were cultured using Dulbecco’s modified Eagle’s medium (DMEM; Sigma) with 10% fetal bovine serum (FBS) and 6 mM glutamine (called cDMEM). The TBC9^AID^ parasites were generated by transfecting ∼20 μg of linearized plasmid containing ∼1000 bp of targeting sequence in frame with the 3xHA and AID tags in the RH*Δku80Δhxgprt* strain expressing OsTIR1 driven by the gra1 promoter (59). Recombined parasites were selected using 25 μg/mL mycophenolic acid and 50 μg/mL xanthine (MPA/Xanthine) in the culture medium. Clonal parasite lines were obtained by infecting a 96 well plate with a single parasite per well. The clonal lines were screened for HA signal by immunofluorescence assay (IFA) using the anti-HA antibody (3F10, Roche/Sigma). For CRISPR based transfections, 5 μg of a plasmid expressing a Cas9 gene and a gRNA targeting specific locus was mixed with 50 uL of a Q5 polymerase (NEB) PCR product containing the expression cassette of the gene of interest and a selection cassette flanked by homologous targeting region. The transfectants were initially selected with MPA/Xanthine for 36 hours (CRISPR plasmid selection) and finally with either cDMEM containing 20 μM chloramphenicol or treatment with 40 μg/mL Zeocine in Hanks buffered saline solution (HBSS; Sigma). Clonal lines were obtained in a similar manner to the TBC9^AID^ parasite line. The mRuby3-Ran reporter strain was generated by CRISPR-mediated targeting of the endogenous ATG start codon of Ran (TGGT1_248340), and replacing the coding sequence with a mRuby3-Ran open reading frame followed by a downstream chloramphenicol selection cassette. The nucleocytoplasmic reporter stain was generated by CRISPR targeting a region of the empty Ku80 locus by and inserting a chloramphenicol resistance cassette and the TGGT1_258850^3xFLAG^ ORF driven by the gra1 promoter. Both the mRuby3-Ran and NCT reporter strains were then modified to tag the TBC9 with AID-3xHA, as above. Additional wild-type or mutant copies of TBC9 were added to these strains by either reporter TBC9^AID^ strain with a CRISPR targeting a second region of the Ku80 locus and inserting a bleomycin resistance cassette and either TBC9-3xTy1 ORF driven by a dhfr promoter.

### Plaque assay

A six well plate containing confluent HFF monolayers was infected with 100 parasites per well from a syringe released T25 of a respective clonal population in cDMEM containing auxin (IAA; 500 μg/mL in 100% ethanol) or control (equal volume of 100% ethanol). The HFF monolayer was fixed with methanol after 8 to 11 days and stained with crystal violet. All plaque assays were performed in n=3 technical replicates for each of the n=3 biological replicates. Plaques were counted using Fiji (60) and the p-values were calculated by unpaired two-tailed Student’s t-test.

### Western blots

Parasite lysates were prepared by pouring hot 1× SDS PAGE loading dye over highly infected HFF monolayer and collecting them by scraping (100 μL/well in a 24 well plate). The lysate was then sonicated, heated at 95°C for 10 min. and spun at 21000xg for 10 min. Cleared lysates were loaded on a 12% SDS-PAGE gel and ran for 200 V for 45 minutes to resolve the proteins. They were transferred to a PVDF membrane at 200 mA for 2 hours in wet transfer system (BioRad). Membranes were blocked for 1 hour in 5% milk in TBST (10 mM Tris-HCl pH 7.5, 150 mM NaCl, 0.1% Tween 20) or 5% BSA in TBST (for anti-Ty1 blot). The blot was incubated in primary antibody overnight at 4°C. Primary antibody dilutions – 1:1000 Rabbit anti-HA in TBS (10 mM Tris-HCl pH 7.5, 150 mM NaCl); 1:20000 anti-Ty1 (MA5-23513; Thermo scientific) in 5% BSA in TBST; 1:1000 rabbit anti-TgTub and 1:1000 rabbit anti-HsRanGAP1 (Cell Signaling Technology #36067; gift of Fontoura lab) in milk blocking buffer. The next day, the blot were washed 3× with TBST to remove primary antibody. The blot was then incubated with either an anti-mouse (BioRad #170-6516) or an anti-Rabbit (Invitrogen #31460) HRP conjugated secondary antibody at 1:15000 in milk blocking buffer and incubated at room temperature for 1 hour. The blot was washed 3× with TBST and imaged on the GE healthcare ImageQuant LAS4000 using ECL Plus reagent (Pierce). The images were processed using Fiji (60).

### Immunofluorescence assay and microscopy

HFFs were grown in a 24 well plate containing glass coverslips until a confluent monolayer formed which was then infected with parasites. Post incubation, the coverslips were washed with 1x phosphate buffered saline (PBS) and fixed with 4% paraformaldehyde and 4% sucrose in PBS at room temperature for 15 min. The coverslips were washed with PBS 3× and then permeabilized with 0.1% Triton X-100 in PBS for 30 min. 3% Bovine Serum Albumin (BSA; Sigma) in PBS was used as blocking buffer and incubated for 1 hour. The coverslip was moved to a humidity chamber and respective mix of primary antibodies (diluted in blocking buffer) were added and incubated overnight at 4°C. The next day, the coverslips were washed with permeabilization buffer (0.1% Triton X-100 in PBS) 3× with 5 min incubation between each wash. A mix of secondary antibodies conjugated with Alexa-flour and diluted in blocking buffer was added to the coverslips and they were incubated for 1 hour at room temperature. Three washes with permeabilization buffer were performed to remove the secondary antibody and Hoechst stain (in PBS) was added and incubated for 15 min. The coverslips were washed with PBS and then mounted on a glass slide using mounting medium (Vector Laboratories). The cells were imaged on a Nikon A1 laser point scanning confocal microscope with a 60× oil immersion 1.42 NA objective using Nikon Elements. Primary antibodies used in this study are rat anti-HA (1:1,000 dilution; Sigma #11867423001), rabbit anti-Tg-β-tubulin (1:10,000 dilution), rabbit anti-TgSec61 (1:1000; preconjugated to AlexFlour 647) mouse anti-Ty1 (1:20,000; #MA5-23513 Invitrogen), and mouse anti-FLAG (1:1000; Sigma #F1804). The secondary antibodies used in the study are Goat anti-Rabbit IgG, AlexaFluor 647 (ThermoFisher; #A21245), Goat anti-Mouse IgG, AlexaFluor 555 (ThermoFisher #A21424) and Goat anti-Rat IgG, AlexaFluor 488 (ThermoFisher #A21208).

### Phenotype quantification

Clonal population of either mRuby3-Ran or NCT reporter containing either TBC9^AID^ only or with the rescue copies of TBC9 were syringe released and counted on a hemocytometer. 0.7 million parasites per well were added to HFF monolayer in a 24 well plate containing coverslips. The plate was incubated for 4 hours and then washed with 1 mL cDMEM twice to remove uninfected parasites. The media was changed to cDMEM containing auxin (IAA; 500 μg/mL in 100% ethanol) or control (equal volume of 100% ethanol) for 2 hours. The coverslips were fixed at the end of 2 hours and IFA stained with Hoechst, anti-HA, anti-Tub or anti-Ty1 for mRuby3-Ran, and Hoechst, anti-HA, anti-FLAG and anti-Tub for the NCT reporter parasites. The parasites were imaged on the Nikon A1 pscm with a 60x oil immersion objective with a 1.42 NA. The four channel (405nm, 488 nm, 561 nm and 640 nm) Z-stacks were taken with a step size of 0.5 µm, pinhole of 1 AU at TRITC (561 nm) and a resolution of 19.3087 pixels per micron (1024×1024). These settings were maintained across all acquisitions for all images used for phenotype quantification. The stacks were opened in Fiji (60) and the background was subtracted with a rolling ball radius of 40 pixels. Hoechst signal was used to mark the nucleus. mRuby3 signal in the mRuby3-Ran and tubulin signal in the NCT reporter parasites were used to mark the parasites. The average intensity of nuclear signal was divided by the average intensity of the corresponding parasite to get nuclear/total mean intensity for each parasite. These value were used for comparing the ± auxin conditions graphically and for calculation of average values to finally compare them using an unpaired Student’s t-test for the TBC9^AID^ parasites and the parasites with the rescue copies of TBC9.

### Recombinant protein purification

All protein cDNAs were cloned in a pET28a or pGEX-4T-1 base vector with either a His6, or His6-SUMO, or GST-His6 tag. All proteins were expressed in the *E. coli* Rosetta 2 cells (Novagen, Sigma) and induced with 200 µM isopropyl β-ᴅ-1-thiogalactopyranoside (IPTG) between the OD_600nm_ of 0.6 – 0.8, and incubated for 20 hours at 16°C. Cells were harvested and stored until the day of purification. His6-TgRan and His6-TgRan^Q70L^, and His6-TgRan^Q70L/R96A/K100A/R107A^ (basic patch mutant; TgRan^R/K-Mut^) used for GTPase assay, were purified with Ni-NTA affinity chromatography followed by anion exchange chromatography (GE Healthcare HiTrap Q HP column; Buffer A: 10 mM Tris HCl pH 8.0, 5 mM MgCl_2_; Buffer B: Buffer A + 1M NaCl). The RanGTP peak (24 to 32 %B) was concentrated and frozen in aliquots for the GTPase assay. The GST-His6-TgRan, from here on annotated as GST-TgRan, was purified with Ni-NTA affinity chromatography, anion exchange chromatography followed by a size exclusion chromatography (SEC) in the GE healthcare Superdex 200 16/60 pg column in 10 mM HEPES pH 7.5, 150 mM NaCl, 5 mM MgCl_2_. Final protein was concentrated and stored until further use. His6-SUMO-TgRan was expressed and purified through anion exchange. The eluted protein was ULP1-digested to remove the His6-SUMO and dialyzed against 10 mM Tris HCl pH 8.0, 25 mM NaCl, 20 mM MgCl_2_. The protein was then applied to the HiTrap Q HP column with Buffer A: 10 mM Tris HCl pH 8.0, 20 mM MgCl_2_; Buffer B: Buffer A + 1M NaCl). A majority of untagged TgRan came out in the flowthrough which was SEC purified as above. The Rab1B(1-180), Rab2(1-188), Rab11A(1-186), and Rab11B(1-189) and their Q to L mutants (Rab1B Q67L, Rab2 Q66L, Rab11A Q71L, and Rab11B Q71L) were expressed with a His6-SUMO tag, Ni-NTA affinity purified, cleaved the tags with ULP1, anion exchange purified and GTP containing Rab fraction was frozen. The human RanGAP (1-406) and human RCC1 were expressed with a His6-SUMO tag and the tags were cleaved with ULP1 after Ni-NTA purification. Both the TBC9 variants, the human RanGAP (residues 1-406) and human RCC1 were further purified by anion exchange and SEC as used for GST-TgRan. The human polynucleotide phosphorylase (hPNP; gift of Xuewu Zhang) was expressed as a His6 tagged protein and purified with the same three step protein purification method.

### GTPase assays

To track the TBC9 RanGAP activity in lysates we used the GTPase Glo assay (Promega) (33), modified as follows. GST-TgRan was preloaded with GTP and assayed for single-turnover hydrolysis. To load the protein with GTP, GST-TgRan was bound to glutathione magnetic resin (#78602; Pierce) and incubated with 20 mM EDTA and 1 µM recombinantly purified human RCC1. MgCl_2_ was added after 2 hours to a final concentration of 40 mM to saturate the EDTA. The GST-TgRan protein was eluted with 20 mM Glutathione in 10 mM HEPES and 150 mM NaCl after three washes with 10 mM HEPES and 150 mM NaCl. The protein concentration was measured by absorption at UV_280nm_. 5 µL of 10 µM GTPase was used per GTPase Glo reaction. After the incubation unhydrolyzed GTP was read-out by luminescence per kit instructions.

The kinetic GTPase assays were adapted from (50, 61, 62) and were performed at 24°C with 20 µM GTP loaded His6-TgRan or His6-TgRan^Q70L^ protein, 150 μM 7-Methyl-6-thioguanosine (MESG; Berry & Associates Inc) and 2.91 μM (0.1 μg) of recombinantly purified human phospho-nucleotide phosphorylase (PNP) in a 100 uL reaction with assay buffer (10 mM HEPES pH 7.5, 100 mM NaCl, 5 mM MgCl_2_). In case of Rabs, we used 80 µM of GTPase for each reaction. 5 uL of respective effectors at varied concentrations were added to initiate the reaction, which was monitored by measuring change in absorbance at 360 nm over time in a plastic UV cuvette (BRANDTECH Scientific, Inc.) with a 1 cm path length. We observed a small nonspecific linear drift in OD_360_ in all reactions containing TBC9. This was subtracted as background before further analysis. To obtain *k_obs_*, the kinetic data were fit to either a linear (pseudo-zeroth order) or inverse exponential (pseudo-first order; Abs(*t*)=( *A* − *A*)⋅(1−e^−^*^kobs^*^⋅^*^t^*)+ *A*_min_). To obtain *k* / *K* , the calculated *k*_obs_ values from first-order fits were plotted against the concentration of TBC9 used in each reaction and fit to the linear equation *k_obs_*=(*k_cat_* / *K_m_*)⋅[*TBC* 9]+*k* _int_, where *k* _int_ is the intrinsic activity of the GTPase.

To quantify the nucleotide content of the proteins, bound nucleotides were extracted using the methanol-chloroform protein extraction method (63). Instead of the protein layer from centrifugation step, we took the aqueous layer containing the nucleotides and applied it to the Capto Hires Q 5/50 column (Cytivia). Nucleotides were eluted with a gradient of 15-22% buffer B (peaks: GDP ∼18% buffer B; GTP ∼20% buffer B). The chromatogram was plotted matplotlib and peaks integrated using scipy.

### Analytical size exclusion chromatography

To form complex, 65 μM TgRan was mixed with 195 μM TgTBC9 protein in presence of 65 μM GDP, 2 mM AlCl_3_ and 20 mM NaF, and incubated on ice for 2 hours. The complex was loaded on a Superdex 200 Increase 10/30 (Cytivia) gel filtration column pre-equilibrated with 20 mM Tris-pH 7.5, 100 mM NaCl, 2 mM MgCl_2_, 2 mM DTT, 20 mM NaF. The individual proteins were separated by gel filtration as above, without pre-incubation and without AlCl_3_ or NaF. Equal volumes of each fraction were loaded on a 12% SDS-PAGE gel and visualized with Coomassie stain.

### TBC9 enrichment, mass spectrometry, and identification from parasite lysate

20× 15 cm dishes were infected with ∼20 million parasites each, incubated for ∼36 hours and mechanically disrupted with a 21 gauge needle. The freed parasites were filtered through a 5 µm filter to remove host cell debris (and human RanGAP contamination), and centrifuged to pellet the parasites. All following steps were conducted on ice or at 4°C, as appropriate. The parasites were resuspended in 12 mL of Lysis buffer (20 mM Tris-HCl pH 8.0, 100 mM NaCl, 1 mM PMSF, 1x Roche Protease inhibitor cocktail) and lysed by sonication using a medium probe on a Branson 450 Digital Sonifier. The lysate was ultra centrifuged at 100k×g for 50 min in a TL100.3 rotor. The supernatant was collected in a fresh 50 mL tube and saturated ammonium sulfate was added slowly, with continuous mixing, to a final concentration of 40% at 4°C in a cold room. The lysate was stirred for 30 min to facilitate precipitation. The turbid solution was ultracentrifuged at 100k×g for 30 min. The unprecipitated protein was was dialyzed against 10L of 20 mM Tris-HCl pH 8.0, 100 mM NaCl for 6 hours with 4 buffer changes. The dialyzed lysate was taken in a fresh 50 ml tube and applied to a 1 mL HiTrapQ column equilibrated with buffer A (20 mM Tris HCl pH 8.0) without further dilution. The proteins bound to the column were eluted with a gradient of 15-50% buffer B (20 mM Tris HCl pH 8.0, 1 M NaCl) and 0.5 mL of fractions were collected. All fractions were screened for RanGAP activity using the GTPase Glo kit, and specific activity was calculated as the highest dilution at which activity could still be identified (see below). 10 Fractions, corresponded to 37 to 46 % buffer B, containing the highest RanGAP activity were sent for tandem mass tag (TMT) protein identification mass spectrometry. To control for potential errors in the sequence of fractions during sample preparation for mass spectrometry, we doped them with increasing amounts (1 to 10 ng) of recombinantly purified *Neospora canninum* ROP4. In addition, we added 5 ng of mouse IRG6a was added to each fraction to assist in normalization of total protein abundance.

For TMT mass spectrometry sample preparation, 30 uL of 5% SDS was added to each sample to bring the total volume to 50 uL. Following disulfide bond reduction and alkylation with tris(2-carboxyethyl)phosphine (Sigma-Aldrich) and iodoacetamide (Sigma-Aldrich), samples were digested overnight with trypsin using S-Trap micro columns (Protifi). The peptide eluates from the S-Traps were dried and reconstituted in 50 uL of triethylammonium bicarbonate (TEAB) buffer. The TMT 10plex Isobaric Mass Tagging Kit (Thermo) was used to label the samples as per the manufacturer’s instructions, quenched with 5% hydroxylamine, and the samples were combined and underwent solid-phase extraction cleanup using an HLB elution plate (Waters). The samples were then dried in a SpeedVac and reconstituted in a 2% acetonitrile, 0.1% TFA buffer and diluted such that ∼1 ug of peptides was injected. Peptides were analyzed on a Thermo Orbitrap Eclipse MS system coupled to an Ultimate 3000 RSLC-Nano liquid chromatography system. Samples were injected onto a 75 um i.d., 75-cm long EasySpray column (Thermo) and eluted with a gradient from 0-28% buffer B over 180 min at a flow rate of 250 nL/min. Buffer A contained 2% (v/v) acetonitrile and 0.1% formic acid in water, and buffer B contained 80% (v/v) acetonitrile, 10% (v/v) trifluoroethanol, and 0.1% formic acid in water. at a flow rate of 250 nl/min. Spectra were continuously acquired in a data-dependent manner throughout the gradient, acquiring a full scan in the Orbitrap (at 120,000 resolution with a standard AGC target) followed by MS/MS scans on the most abundant ions in 2.5 s in the ion trap (turbo scan type with an intensity threshold of 5,000, CID collision energy of 35%, standard AGC target, maximum injection time of 35 ms and isolation width of 0.7 m/z). Charge states from 2-6 were included. Dynamic exclusion was enabled with a repeat count of 1, an exclusion duration of 25 s and an exclusion mass width of ± 10 ppm. Real-time search was used for selection of peaks for SPS-MS3 analysis, with the search performed against the human reviewed protein database from UniProt and a custom protein database containing *Neospora canninum* ROP4 and mouse IRG6a sequences in addition to all *Toxoplasma* protein sequences. Up to 1 missed tryptic cleavage was allowed, with carbamidomethylation (+57.0215) of cysteine and TMT reagent (+229.1629) of lysine and peptide N-termini used as static modifications and oxidation (+15.9949) of methionine used as a variable modification. MS3 data were collected for up to 10 MS2 peaks which matched to fragments from the real-time peptide search identification, in the Orbitrap at a resolution of 50,000, HCD collision energy of 65% and a scan range of 100–500. Protein identification and quantification were done using Proteome Discoverer v.3.0 SP1 (Thermo). Raw MS data files were analyzed against the above database. Both Comet and SequestHT with INFERYS Rescoring were used, with carbamidomethylation (+57.0215) of cysteine and TMT reagent (+229.1629) of lysine and peptide N-termini used as static modifications and oxidation (+15.9949) of methionine used as a variable modification. The Reporter Ions Quantifier node within Proteome Discoverer was used for quantitation, using reporter ion intensities. The false-discovery rate (FDR) cutoff was 1% for all peptides.

The final protein identification data were then normalized using IRG6a abundance and the sequence of fractions verified with increasing amount of NcROP4. The abundance of all identified proteins were correlated with the RanGAP activity across each sample by calculating Pearon’s correlation coefficient in SciPy. Since the Promega GTPase assay is semi-quantitative, we measured the highest dilution of each fraction that exhibited RanGAP activity higher than 40% the activity of a 5 µM human RanGAP (1-406) control in a 60 min reaction.

### Yeast RanGAP complementation

The base temperature-sensitive RanGAP *rna1-1* strain was created by transforming w303 *S. cerevisiae* by targeting the rna1 locus with a Q5 PCR product encoding the NatMX selection cassette and the *rna1-1* mutations (36, 64). Recombinant yeast were selected with nourserothricin and replica plated to identify colonies which exhibited temperature-sensitive growth. To test for rescue of RanGAP function in yeast, the *rna1-1* w303 strain was transformed with a plasmid containing the ORF for the gene of interest driven by the constitutive ADH1 promoter and a Leu2 selection cassette. Transformants were selected by growth in -Leu media. Rescue of RanGAP function was assessed by comparing growth in permissive (24°C) versus restrictive (37°C) temperatures.

### Bioinformatic analysis

The sequence of yeast, *Toxoplasma*, and the red algae *Chondrus crispus* LRR*-*containing proteins (InterPro IPR001611) were taken from ToxoDBv68 and UniProt. These sequences were combined with the RanGAP sequences from human, *Drosophila*, *S. pombe*, *Xenopus laevis*, *Arabidopsis thaliana*, the Stramenopile *Saprolegnia parasitica*, the Rhizaria *Plasmodiophora bassicae*, and the ciliate *Tetrahymena thermophila.* LRR sequences were aligned with MAFFT (65). To assess TBC phylogeny, sequences of all TBC domains (PFAM PF00566) from each of the organisms used in a previous analysis of the evolution of TBC domains (39) were combined with the hits from a PSI-BLAST search to convergence using TgTBC9 as query with a 1e-50 cutoff. The aligned Alveolate TBC9 domains were used to perform a search with HMMERv3.4 against the core SAR and haptophyte proteomes from EukProtV3 (66). The top hit from each organism was added to the list of unaligned sequences. All TBC sequences were aligned with ClustalO (67) using the aligned hits from the PSIBLAST search as a seed profile. Phylogenetic trees were estimated by IQTREE2 using SH-aLRT (68) and ultra-fast bootstrapping (69) and the model selected automatically (support values are reported as “SH-aLRT/bootstrap” in the resulting trees). Accessions and phylogenetic trees are in SI Data S1,4,7. Alignments are in SI Data S2-3,6,8. Organism phylogeny in Figure S8 was downloaded from the Open Tree of Life (70). The phylogenetic trees were visualized using Archaeopteryx from Forester (71) and further annotated in Inkscape. Alphafold3 server (72) was used for generating models of the TgRan:TgTBC9 complex, which were analyzed in PyMOL (73).

### Figure generation

All figures were generated in Inkscape v1.4. Graphs were generated using matplotlib and seaborn packages from python. Statistical analyses were conducted in python with the SciPy and Statsmodels packages.

## Data Availability

Proteomics data from SI Data S5 are available on ProteomeXchange accession PXD077025.

## Supporting information

SI Data 1 - Accessions for phylogenies

SI Data 2 - LRR alignment for Figure S1

SI Data 3 - extended LRR alignment

SI Data 4 - extended LRR phylogeny

SI Data 5 - Mass spectrometry data

SI Data 6 - TBC full alignment

SI Data 7 - TBC full phylogeny

SI Data 8 - TBC9 subalignment

SI Data 9 - List of primers

## Acknowledgments

We acknowledge the Proteomics Core Facility at University of Texas Southwestern Medical Center for mass spectrometry analysis. We thank Elliott Ross and Joseph Albanesi for discussions on GTPase biochemistry, Boyuan Wang and members of the Xuewu Zhang lab at UTSW for advice regarding GTPase assays, Jonathan Friedman and Mike Henne for advice on yeast genetic manipulation, and Matt Daugherty for advice on evolutionary analysis. M.L.R. acknowledges funding from the Welch Foundation (I-2075-20240404) and a Burroughs Wellcome Foundation Investigators in Pathogenesis of Infectious Disease award (G-1021959).

---

**Figure S1:**
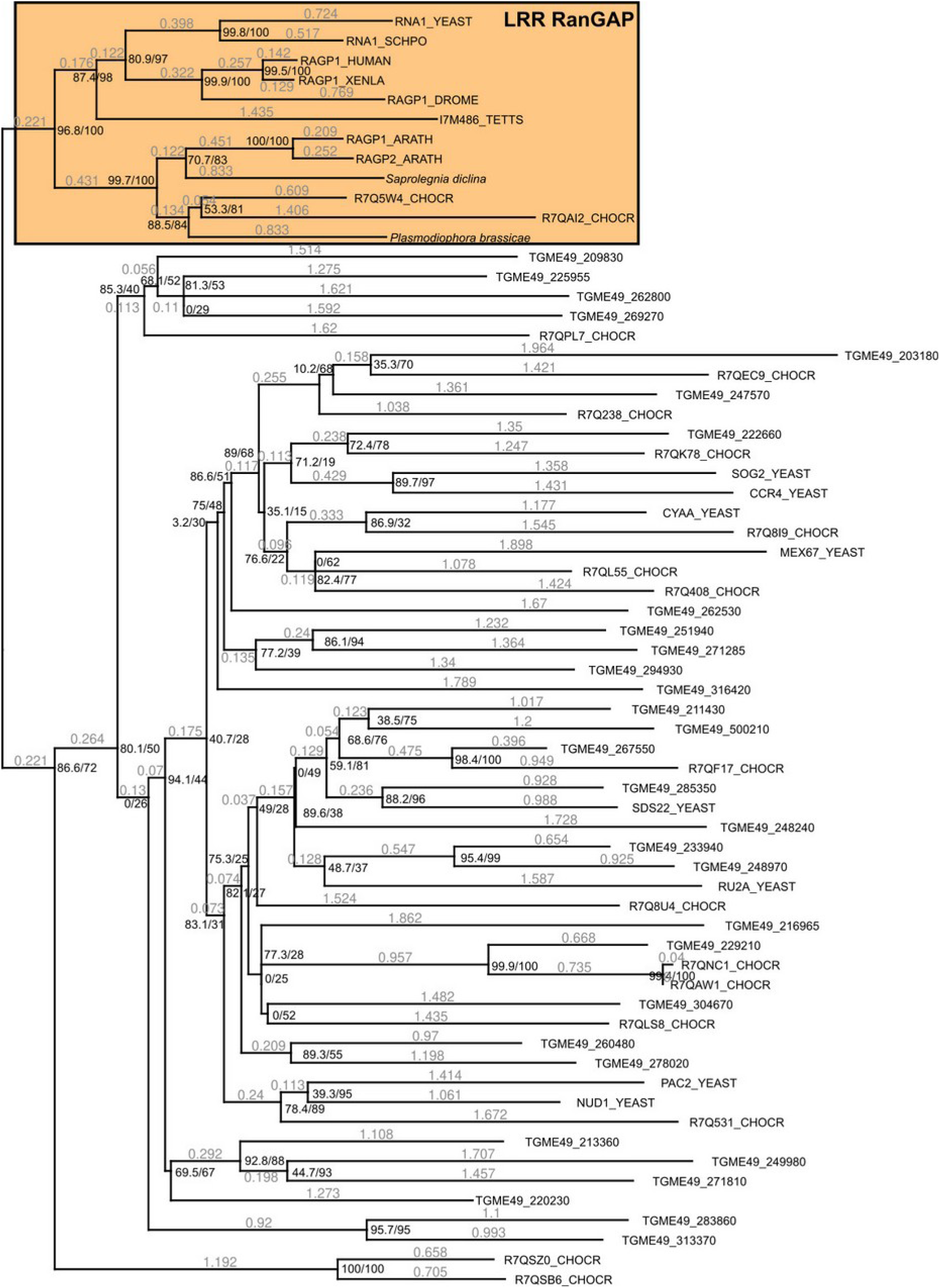
Phylogenetic analysis of *Toxoplasma* LRR-containing proteins. All LRR domains from *Toxoplasma,* baker’s yeast, and *Chondrus crispus* were aligned, together with diverse RanGAPs from opisthokonts, plants, the ciliate *Tetraymena* (an alveolate)*, Saprolegnia* (Stramenopile) and *Plasmodiophora* (Rhizaria). Phylogenetic trees were estimated using IQTREE2 with aLRT/ultrafast bootstrapping (black). Branch lengths are gray (substitutions per site). The RanGAP proteins form a distinct clade from all other proteins, boxed in gold. Protein accessions and an extended phylogeny are available in SI Data S1 & S4. To create the extended phylogeny, first the top hits (lowest e-value) from the core SAR and haptophyte sequences from the EukProtv3 database of a search with the Panther PNTHR24113 RanGAP HMM profile using HMMERv3.4 were aligned with the sequences used to create this figure. Phylogenetic trees were estimated by IQTREE2 using ultra-fast bootstrapping and the model selected automatically.

**Figure S2:**
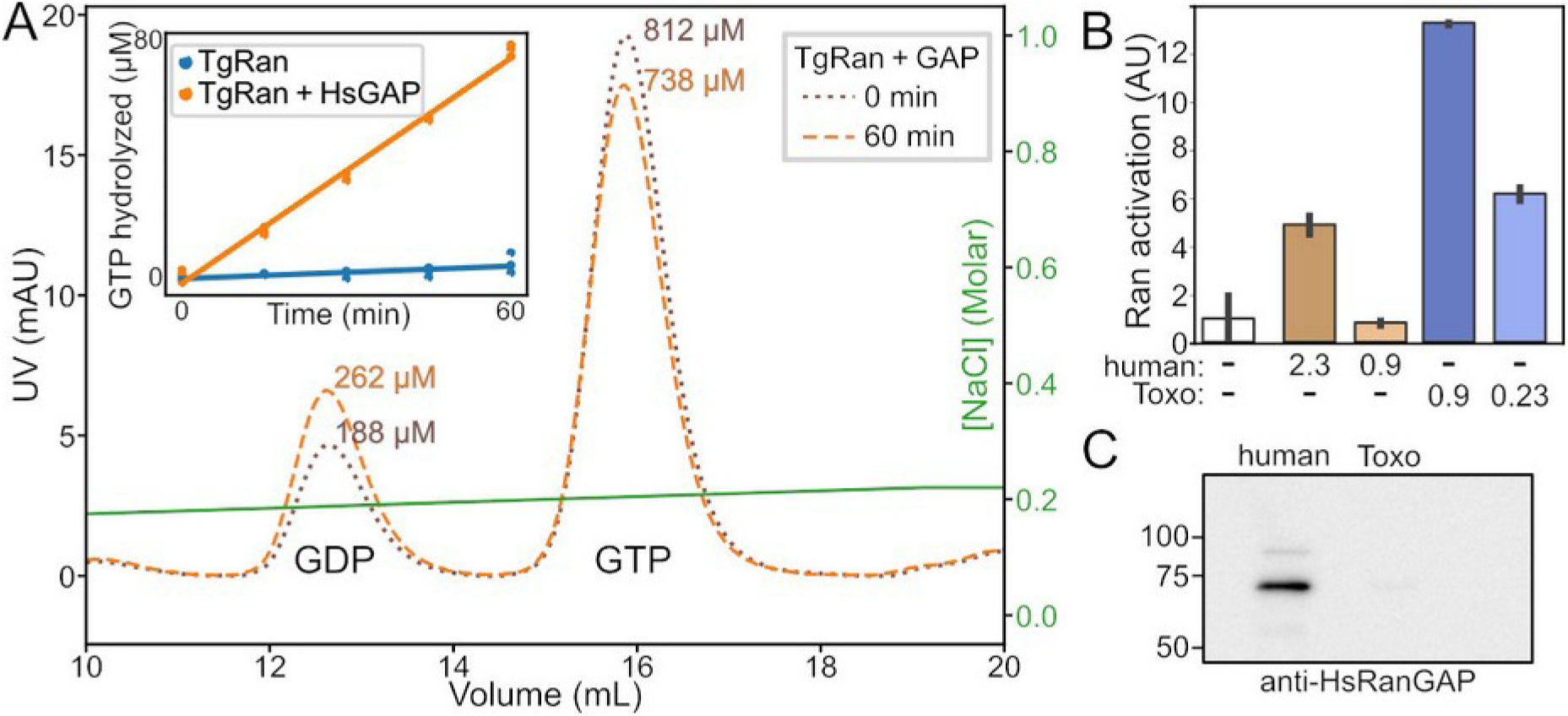
*Toxoplasma* lysate contains RanGAP activity. (A) We used anion exchange chromatography to separate nucleotide from recombinant TgRan that had been incubated at 37°C with human RanGEF and ±HsRanGAP (multi-turnover reaction). Representative chromatograms comparing the 0 and 60 min timepoints are shown, and the entire time course plotted in the inset. (B) Single-turnover reactions using GTP-bound TgRan and the indicated amounts in μg of either human or *Toxoplasma* lysates. Units of activity are the maximum dilution-factor of sample with measurable activity, as described in methods. (C) Western blot probed with anti-human RanGAP comparing 12 μg of lysates from either human and *Toxoplasma* cells demonstrates little contamination of HsRanGAP in the *Toxoplasma* lysate preparation.

**Figure S3:**
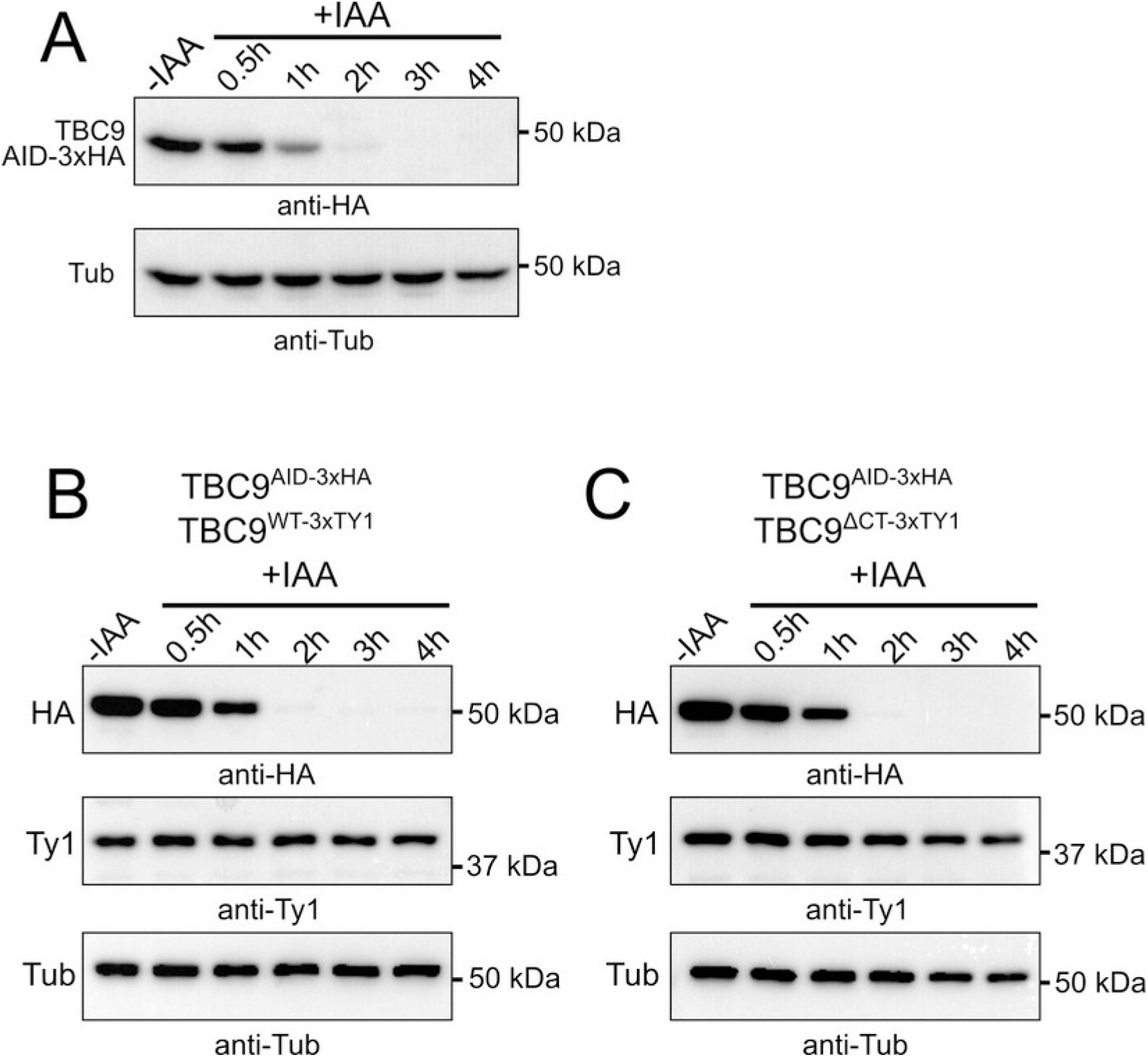
TBC9^AID^ is degraded within 2 hours after auxin treatment. (A) Western blot showing a time course of HA-tagged TBC9^AID^ protein levels during incubation in +IAA media. TBC9^AID^ was probed with anti-HA, and rabbit anti-TgTubβ was used as a loading control. Expression of an additional copy of TBC9 (B) wild-type or (C) mutant does not affect timing of TBC9^AID^ degradation. Blots were probed as in (A) with the addition of an anti-Ty1 antibody to observe the additional expressed copies that had been 3xTy1 tagged.

**Figure S4:**
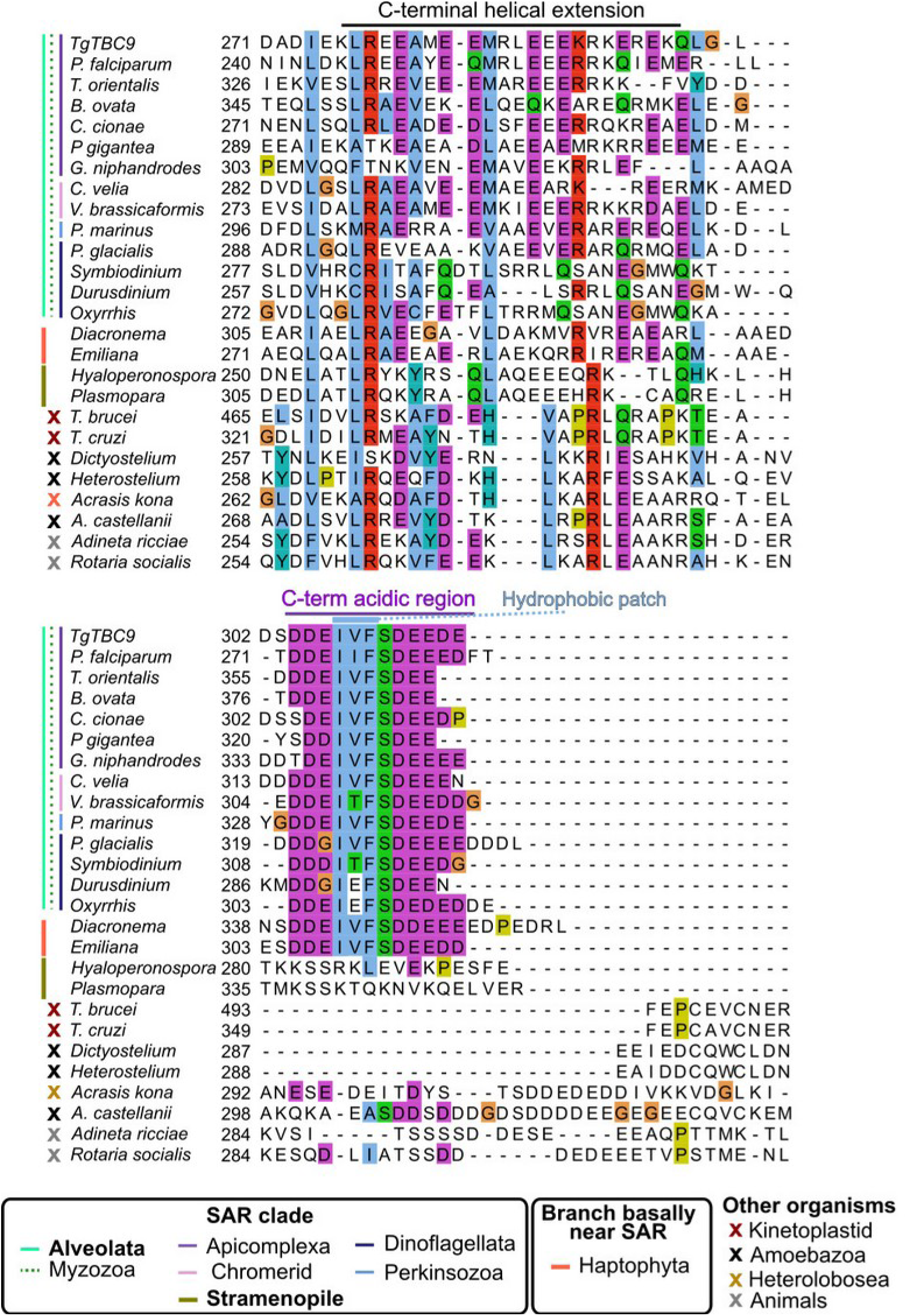
Identification of a conserved C-terminal motif in Alveolate TBC9. Alignment of the last 40∼50 residues of TBC9 (and TBC9-like) domains from the indicated organisms. The conserved extended C-terminal helix and intrinsically disordered acidic low complexity regions are indicated above the aligned sequences. Taxonomic groups are indicated by the colored annotation to the left. Note that the conserved C-terminus distinguishes Myzozoa TBC9 from most other organism’s TBC9-like proteins. The only exception are Haptophytes, which are distantly related to Alveolates. Also note that while some sequences (*e.g.* rotifers) contain acidic stretches, they are not the same distance from the C-terminal helix, nor do they also conserve the central hydrophobic patch found in the Myzozoan TBC9.

**Figure S5:**
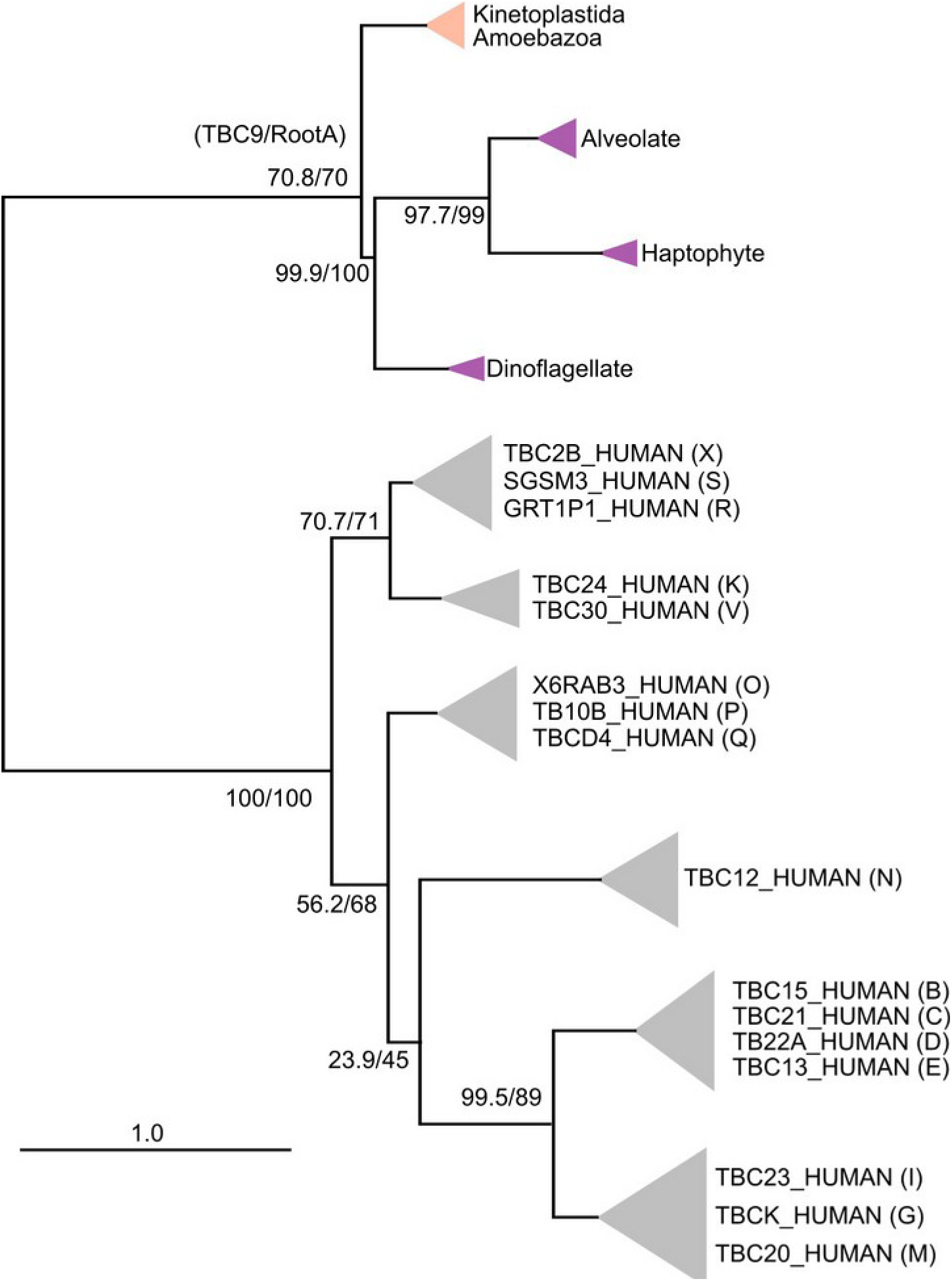
Cartoon of extended TBC phylogenetic tree in SI Data S7. Support values for branches are indicated as SH-aLRT/UF-boostrap. Clades in which all sequences share the *Toxoplasma* C-terminal motif (“TBC9”) are indicated in purple. The “TBC9-like” clade that also belong to the TBC9/RootA family but differ in their C-terminal sequences from *Toxoplasma* is indicated in orange. Previously described TBC families (1) are indicated in parenthesis with representative human sequences. Scale bar indicates substitutions per site.

**Figure S6:**
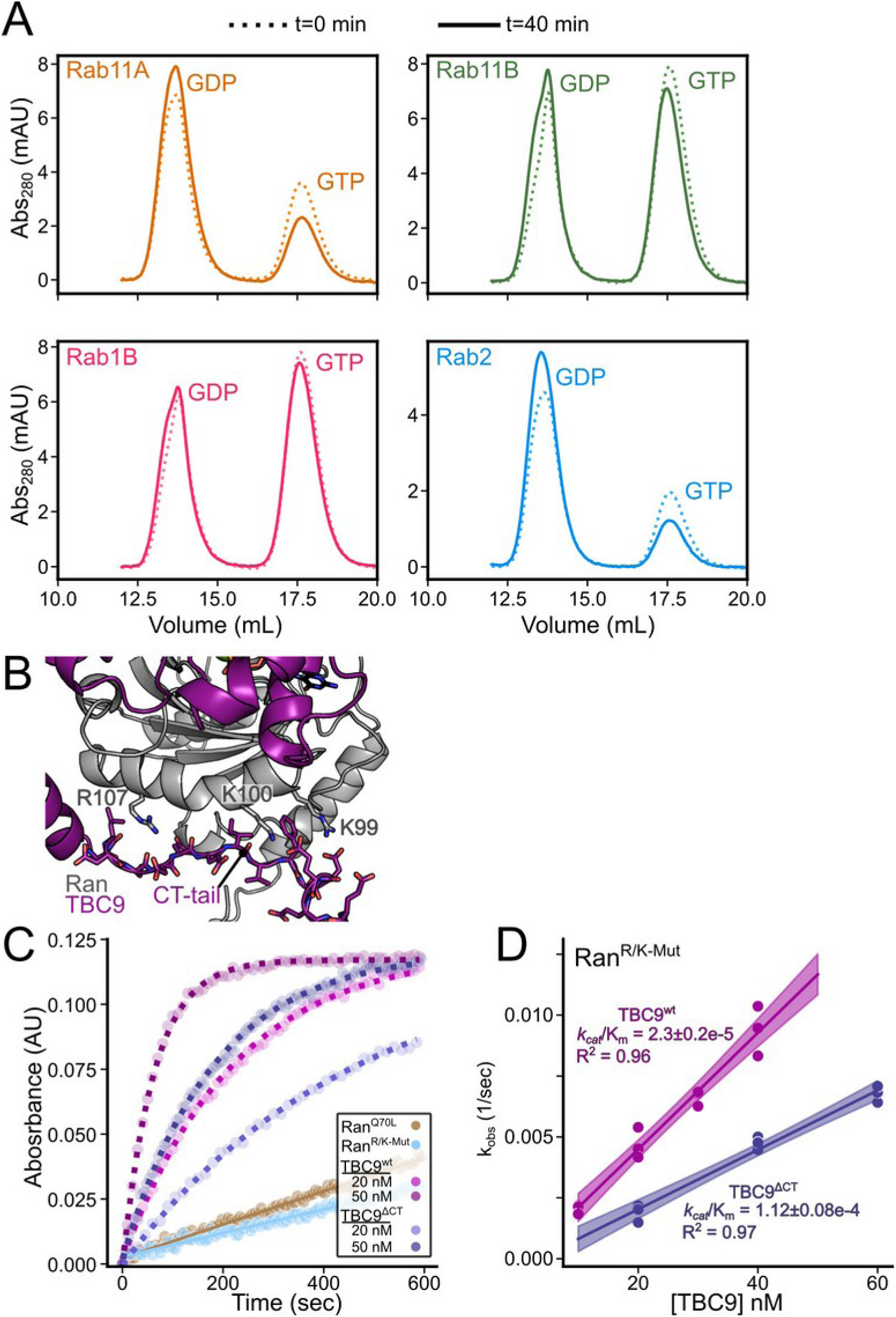
(A) Nucleotide was separated from recombinantly purified *Toxoplasma* Rabs by anion exchange chromatography. All proteins were GTP bound after purification (dotted line), and showed reduced GTP due to intrinsic activity after 40 min incubation at 30°C (solid lines). (B) AlphaFold3 model of Ran (gray) bound to TBC9 (purple), highlighting the predicted interface between Ran and the C-terminal tail. Basic residues in Ran that were mutated are highlighted. (C) The indicated concentrations of TBC9^WT^ or TBC^ΔCT^ were incubated with 20 μM Ran^R/K-Mut^ and the resulting data fit to pseudo-first order kinetics (dotted lines). (D) kobs from fits from (C) graphed versus [TBC9] enables quantitative comparison of TBC9^wt^ versus mutant activities, with error expressed from confidence in fit. Shaded error for all fit curves is 95% ci.

**Figure S7:**
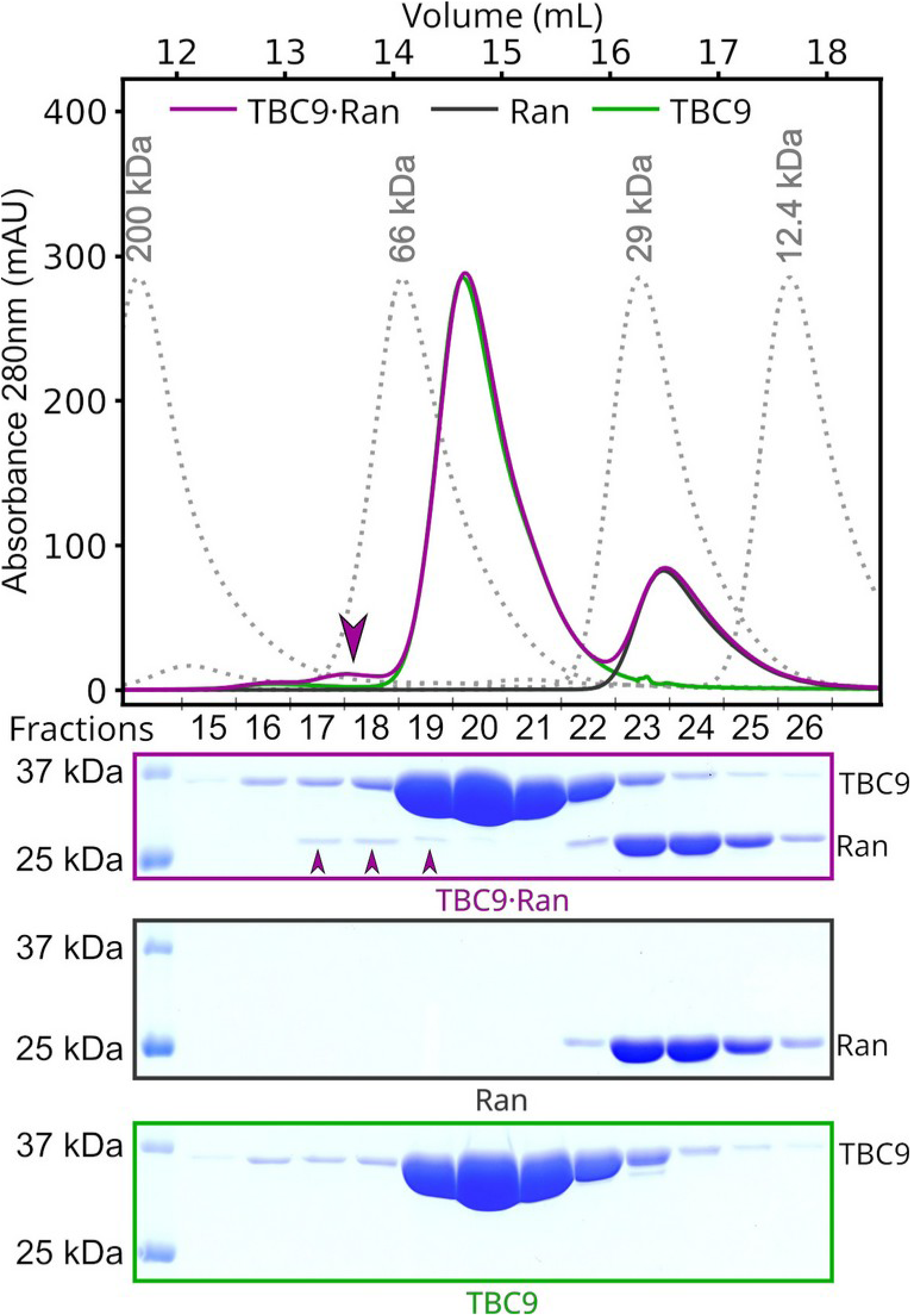
Size exclusion chromatography of 65 μM TgRan (dark gray), 195 μM TBC9 (green) or in complex in the presence of the transition state analog AlF3 (purple). Dotted lines indicate chromatograms from the indicated molecular weight markers. The complex was incubated on ice for 2 hours in the presence of 65 μM GDP, 2 mM AlCl_3_ and 20 mM NaF prior to chromatography. Gels from the resulting fractions from each experiment are shown below. Note that a small amount of Ran (25 kDA) and TBC9 (36 kDa) shift to fractions 17-19 (purple arrows), indicating a heterodimeric complex.

**Figure S8:**
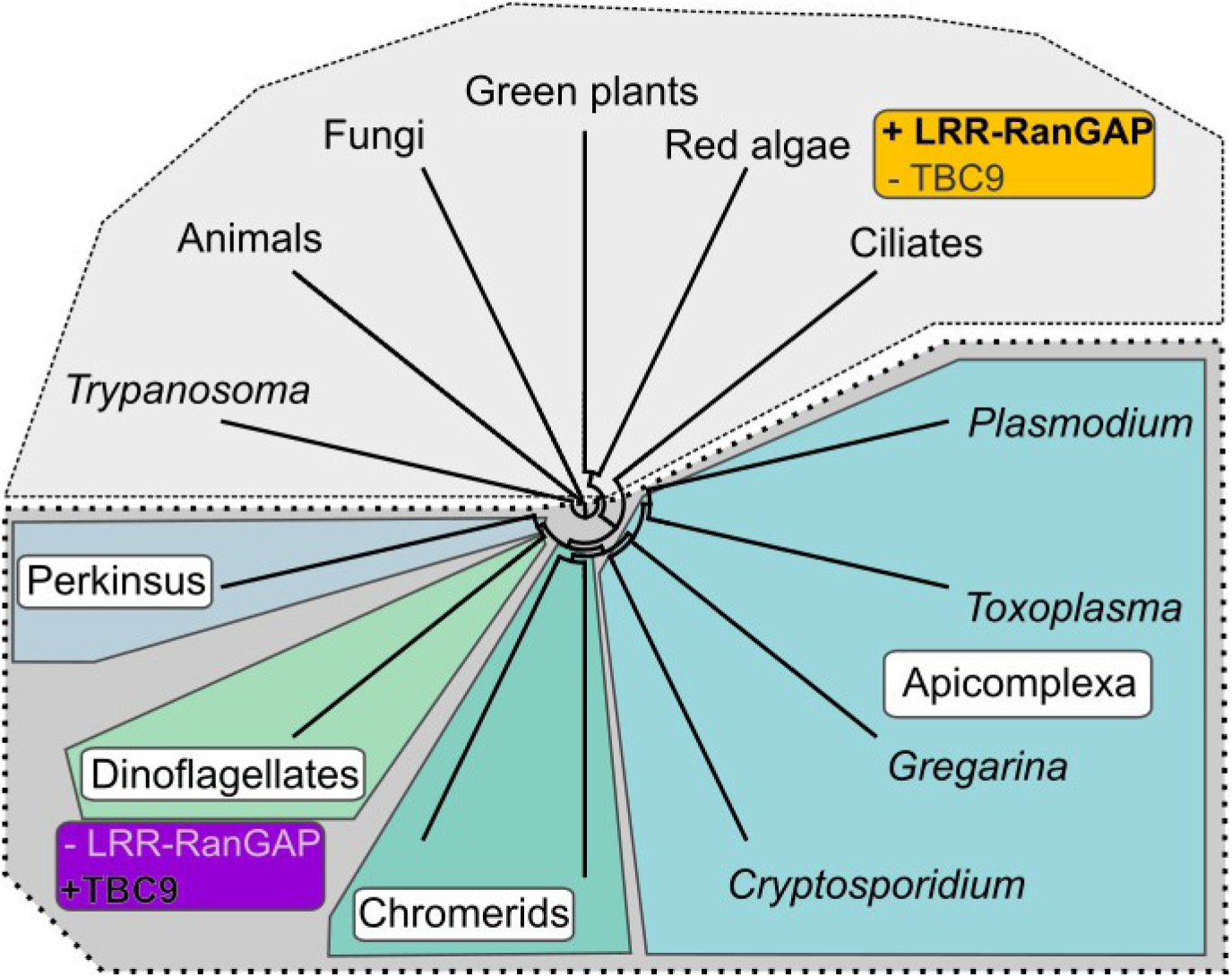
Cladogram representing alveolates and non-alveolate outgroups. Organisms with a typical LRR-RanGAP are shaded light gray. Note all Myzozoa (non-ciliate alveolates) appear to encode a RanGAP TBC9 and are further highlighted in shades of blue and green. Data supporting the presence or absence of an LRR-RanGAP and presence or absence of a TBC9 with the conserved Myzozoan C-terminus are in SI Data S3 (LRR/RanGAP) and SI Data S6 (TBC).

